# Connectivity of restored Atlantic Forest fragments drives composition and functionality of the fungal community in the leaf litter layer

**DOI:** 10.1101/2025.10.14.682367

**Authors:** Guilherme Lucio Martins, Dina in ’t Zandt, Luis Fernando Merloti, Wanderlei Bieluczyk, Gabriel Silvestre Rocha, Robert Timmers, Ricardo Ribeiro Rodrigues, Siu Mui Tsai, Wim H. van der Putten

## Abstract

The restoration of biodiversity and functional tropical forests is critical to mitigating global biodiversity losses. Aboveground, increasing the connectivity of regenerating forests fragments facilitates the recolonization of tropical forest biodiversity. However, restoring functional ecosystems also requires the recovery of decomposition processes as these are essential in shaping aboveground biodiversity. Therefore, we investigate the role of forest connectivity on restoring the composition and functioning of fungal communities in the leaf litter layer during a chronosequence of forest restoration. In the Brazilian Atlantic Forest, we studied secondary forests regrown between 18 to 55 years after deforestation and different levels of forest connectivity and compared their litter to recently abandoned pastures and undisturbed primary forests. We quantified how forest age and connectivity between fragments influenced the litter fungi composition in relation to tree diversity, litter chemistry, and litter isotopes. We show that fungal composition was highly heterogeneous in forest litter, whereas pasture litter exhibited a more homogeneous community. Moreover, forest connectivity had stronger effects on litter fungal composition compared to forest age. Connectivity promoted wood saprotrophs and endophytes, while suppressed soil saprotrophs, with its effects being more evident during later stages of restoration. Fungal guilds such as endophytes and saprophytes, were primarily influenced by tree diversity and leaf litter chemistry. We conclude that forest connectivity promotes the re-establishment of saprophytic fungi capable of decomposing recalcitrant litter substrates, driven mainly by enhancing tree diversity and litter quality. Practical implications of increasing connectivity may relate to forest resilience on front of future climate change scenarios.

## 1. Introduction

The Brazilian Atlantic Forest biome encompasses 1.6 million hectares and stands as one of the global biodiversity hotspots, harboring diverse and endemic flora and fauna and storing an estimated 315 to 460 Mg of carbon per hectare in above- and belowground biomass (Vieira et al. 2011). However, centuries of deforestation driven by urbanization, agriculture, and livestock expansion have fragmented native habitats, reducing connectivity and isolating populations within the remaining forest fragments. This fragmentation limits the movement of fauna and flora, increases the risk of inbreeding and local extinction, and ultimately threatens the functional biodiversity and ecosystem stability (Saura et al. 2017). The current primary forest is fragmented and covers only 11-16% of its original area, and in many cases, is vastly degraded (Rezende et al. 2018; de Lima et al. 2020). Since 1970, efforts to restore Atlantic Forest (Rodrigues et al. 2009) aimed to preserve the biodiversity of primary forests while reconnecting the remaining fragments to support species dispersal, forest regeneration, ecosystem resilience, and climate change mitigation (Poorter et al. 2021; Rother et al. 2023). These efforts have focused largely on converting degraded pasture into secondary forests, as pastures are the main cause of connectivity loss. In this context, connectivity metrics, such as surrounding land use and the percentage of forest cover, provide valuable tools for assessing the regeneration potential of forest fragments (César et al., 2021).

While current restoration efforts focus primarily on restoring aboveground biodiversity, relatively little knowledge exists on microbial responses to forest restoration. For instance, belowground microbes and soil fauna are important drivers of organic matter decomposition and recovery of soil biogeochemical processes (Wang and Kuzyakov, 2024), thereby supporting forest restoration. Yet, these processes have not been examined in the context of different levels of forest connectivity during restoration. Litter decomposition by saprotrophic bacteria and fungi transfers carbon and mineral nutrients from plant litter to the soil foodweb (Baldrian 2017; Liu et al. 2023). Such nutrients become available for plant uptake, facilitating tree growth and triggering a cascade of symbiotic interactions that revitalize forest biodiversity, which may eventually reach levels comparable to those of primary forests (Bardgett and Van Der Putten 2014; Chai et al. 2019). However, there is a major knowledge gap on the relationship among connectivity of forest fragments, leaf litter attributes such as biomass and nutrient concentration, and the composition of fungi in the leaf litter of these fragments. Understanding how forest connectivity influences the composition and functionality of leaf litter fungi may help predict and possibly steer the recovery of forests in restoration programs.

Increasing forest connectivity leads to ecological corridors that enhance the dispersal of aboveground species, thereby increasing both diversity and genetic variability of fauna and flora in the habitat fragments (Salviano et al. 2021). As the restored forests ages, it increases the chance of dispersing propagules (*i.e.*, leaves, fruits, and seeds) by wind and animal, resulting in increasing the fragment size and connectivity (Timmers et al., 2025; Hatfield et al. 2024). This explains why tree diversity may increase with restoration time (Zucchi et al. 2018). As a result, greater plant diversity in restored sites may in turn, increase litter carbon accumulation and provide higher-quality litter substrate to promote microbial activity for organic matter cycling (Zhang et al. 2022). As a result, it is expected that increasing forest connectivity will influence litter composition and fungal composition, thereby affecting the functions performed by the fungi community in the litter layer.

In the present study, we examined the composition and putative functioning of the fungal community present in Brazil’s Atlantic Forest litter layer in relation to time since the start of forest restoration, and connectivity with nearby forest fragments. We quantified how forest connectivity influences litter chemistry, litter biomass and fungal composition. The research was carried out in a chronosequence of restored forests with varying levels of connectivity among forest fragments. The litter was characterized by means of chemical, molecular, and isotopic analyses, while forest connectivity was measured using satellite imagery. We tested the hypotheses that: (*i*) the age and connectivity of forest restoration affects leaf litter chemistry and its fungal composition; (*ii*) older and more connected secondary forests present higher leaf litter biomass accumulation than younger and less connected ones (*iii*) tree diversity is the most important factor mediating the abundance of fungal guilds.

## 2. Material and methods

### 2.1. Study sites and experimental design

The study was conducted in the mesoregion of Piracicaba, São Paulo, Brazil (Figure S1). The region’s climate is humid subtropical (Cfa, Köppen) with a semi-deciduous seasonal tropical forest vegetation characterized by hot and humid summers, and dry winters, with an average annual precipitation of 1273 millimeters (Alvares et al. 2013). In this region, 15 sites were selected to represent a spatial-temporal gradient of forest restoration including variation in both forest age and connectivity (Table 1). We used three pasture areas adjacent to secondary forests, managed like before the forest was restored, to represent the starting point of forest restoration. In contrast, three primary forests, each with at least 100 years old, represented the target of restoration. There were nine secondary forests, categorized into early (18 – 23 years), intermediate (31 – 38 years), and late (43 – 55 years) stages of forest restoration. All secondary forests were previously abandoned pastures and were restored by passive restoration (Brancalion et al. 2016), which consists in allowing natural regrowth without planting seedlings or fertilization, only preventing disturbances from humans and cattle grazing. The forest fragments varied in proximity to other forests patches, representing different levels of connectivity. The geographic coordinates of all sites are provided in the supplementary material (Table S1).

**Table 1.**
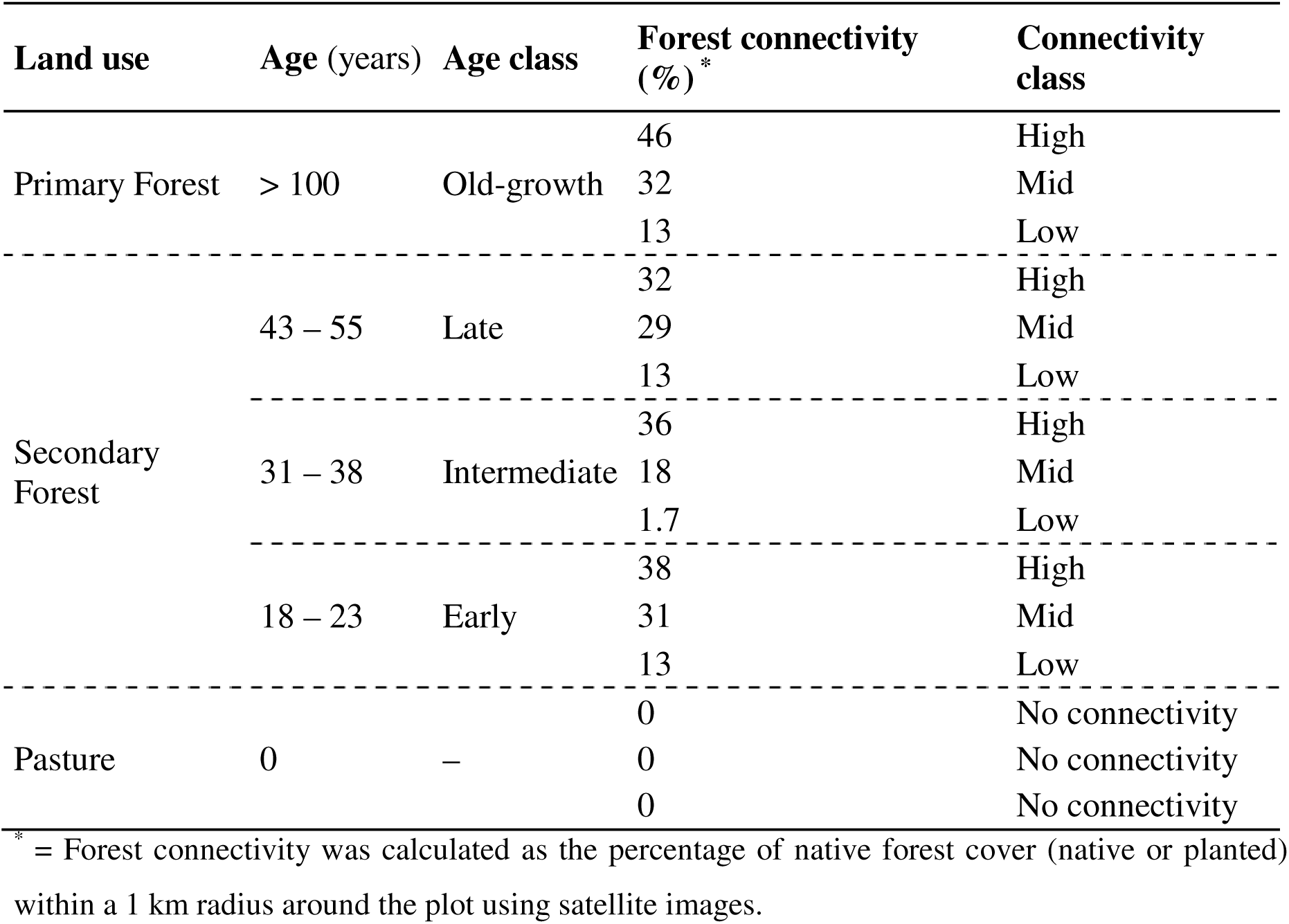
Description of the sites used in this study at the sampling time.

Forest age was provided by researchers who were in charge of the restoration project and confirmed by the stakeholders responsible for each area under restoration. This information was further validated using a chronology of satellite images (MapBiomas – (https://mapbiomas.org/) and historical land cover maps from 1962 to 2008 (César et al. 2021). Forests undergoing restoration were classified into three maturity classes (early/intermediate/late) based on visual indicators such as forest formation type, canopy cover, the presence of late successional tree species, and species with distinct and complementary ecological roles (Rodrigues et al. 2011; Carlucci et al. 2020; Rother et al. 2023).

A proxy of forest connectivity was calculated using high-resolution (5 m) satellite images and MapBiomas forest cover data. Connectivity was defined as the proportion of forest cover within a 1 km radius around each sampling plot, serving as an indicator of habitat availability in the landscape (Rezende et al. 2018; Timmers et al. 2025). Monoculture plantations, such as *Eucalyptus*, were excluded from the calculation, as they do not represent real forest ecosystems. This landscape-scale connectivity metric offers a more robust estimate of ecological connectivity than patch-level metrics such as fragment size or isolation (Fahrig, 2013). Connectivity was categorized into three levels: low (1.7– 13% forest cover), mid (18–32%), and high (33–46%). Each age class included one site of each connectivity level (Table 1).

### 2.2. Forest inventory and litter sampling

Forest inventory was conducted by establishing a 30 × 30 m plot in each site, located at least 50 m away from forest edges and excluding glades and areas impacted by human trails. All trees with a diameter at breast height (DBH) greater than 5 cm were counted and identified to species level (see supplementary information). Plant taxonomic diversity was assessed by the Shannon-Wiener index using the *vegan* package (Oksanen et al., 2019) with the R language.

Litter sampling was conducted in May 2022, at the end of the wet season. For DNA extraction, a mixture of decaying leaf litter was collected from the forest floor, carefully avoiding soil, branches, and wood pieces. Leaf litter from different tree species was not distinguished during sampling. Five samples were collected from each plot (30 × 30 m), one from each corner and one from the center, using 50 mL sterile plastic tubes, each containing approximately 5 g of leaf litter. Samples were then stored and transported in iceboxes to the laboratory for storage in a freezer at -20°C.

For the litter dry biomass weight and chemical analysis, square plastic frames of 25 × 25 cm were used to collect leaf litter material (five replicates per plot). All collected material was carefully taken to exclude soil, branches, and wood pieces. Litter was stored in paper bags and dried at 50°C for 48 hours or until constant weight was achieved.

### 2.3. Litter chemical analysis

Leaf litter samples were ground using a knife mill to achieve a particle size lower than 0.25 mm, ensuring sample homogeneity for element content analysis. Five samples were analyzed per plot. The ground litter was analyzed for total macro (P, K, Ca, Mg, and S) and micronutrient (Fe, Cu, Zn, and Mn) content, according to Malavolta et al. (1989). Briefly, the element concentrations were determined from the extract obtained by digesting the litter material in a mixture of HNO_3_ and HClO_4_. P and S were quantified by colorimetry, while other elements were quantified by atomic absorption spectroscopy (PerkinElmer 3100, USA).

Litter C and N and ^15^N and ^13^C isotopes were analyzed using an elemental analyzer (Carlo Erba, CHN1110; Milan, Italy) coupled to a mass spectrometer (Thermo Fisher, Delta Plus; Bremen, Germany). Stable isotope results were expressed as δ^13^C and δ^15^N (‰) using international standards (Vienna PeeDee Belemnite – V-PDB for C [NBS19 and NBS22] as a reference for ^13^C and composition of atmosphere for N^2^ [IAEA-N1 and IAEA-N2] as a reference for δ^15^N (Farquhar et al. 1982). The delta (δ) values were calculated using the following equation: δ*X* = [(R_sample_/R_standard_) - 1] multiplied by 1000, where *X* refers to ^13^C or ^15^N, and R_sample_ and R_standard_ are the ^13^C/^12^C or ^15^N/^14^N, ratios of sample and standard, respectively.

### 2.4. Litter DNA extraction, sequencing of ITS region, and bioinformatics

DNA was extracted from litter samples using the protocol by England et al. (2004). Briefly, 1.0 g of frozen litter was incubated in 10 mL of sterile 0.5% (w/v) phosphate buffer solution (PBS) at pH 8.0 and shaken for 16 hours at 250 rpm and then centrifuged at 13,000 × g for 20 min at 4°C. The resulting pellet underwent DNA extraction using the DNeasy PowerLyzer PowerSoil Kit (Qiagen, Hilden, Germany) according to the manufacturer’s instructions. DNA integrity was assessed through 1% (w/v) agarose gel electrophoresis, and DNA quantity was measured using a NanoDrop 2000c (Thermo Fisher, Waltham, USA).

A total of 75 samples were sequenced using the fungal internal transcribed spacer (ITS) region with the primers ITS3-2024F/ITS4-2409R (Toju et al. 2012). The library was constructed using the Illumina NextSeq 1000/2000 P1 Reagent kit (Illumina, San Diego, USA), and amplicon fragment size was confirmed by gel electrophoresis. PCR reactions followed the Nextera XT Kit protocol (Illumina, San Diego, USA), and DNA amplification was confirmed by gel electrophoresis. Purified amplicons were equimolarly pooled, and the library’s final concentration was determined using a SYBR green quantitative PCR assay with primers specific to the Illumina adapters Kapa (KAPA Biosystems, Boston, USA).

ITS gene sequencing analysis was performed using the pipeline from *DADA2* (Callahan et al. 2016) package. Briefly, the primers from the demultiplexed data were removed, and data were trimmed and filtered to remove low-quality sequences and, the denoising inference step was performed (based on the learn error step). The forward and reverse sequences were merged (resulting in 2 × 250 bp paired-ended reads), and the chimeric sequences were removed based on the “consensus” method. Amplicon Sequencing Variants (ASVs) were assembled from the retained sequences, and taxonomic assignments were made according to the UNITE database (v. 8) (Nilsson et al. 2019). Data was rarefied to the lowest sample size only for diversity analysis, to reduce overestimation due to differences in sample sizes. The sequences are available at the NCBI Sequence Read Archive under the identification PRJNA1296929.

### 2.5. Statistical analysis

All statistical analyses were conducted in R (version 09.01). To test the first hypothesis, we performed a Principal Component Analysis (PCA) using the *factoextra* package to reduce the dimensionality of litter chemical attributes and compare land-use types. Variables included macronutrients (N, P, K, Ca, Mg, and S), micronutrients (Cu, Zn, and B), and isotopic signature of the litter (δ^13^C and δ^15^N). Fungal composition was transformed into centered log-ratio (*clr*), and the beta diversity was calculated using non-metric multidimensional scaling (NMDS) using the *vegan* package. Differences among land-use types were compared using Permutational Multivariate Analysis of Variance (PERMANOVA) via the *pairwiseAdonis* function. We used Random Forest analysis with the *microeco* package (Liu et al. 2021) to identify fungal biomarker groups within each land-use type. Fungal functional groups were assigned to ecological guilds using the FUNGuild database (Nguyen et al. 2016), considering only classifications marked as “highly probable.”

To test our second hypothesis, we used linear mixed-effects models (LMMs) with the *lme4* package (Bates et al. 2015) to investigate the effects of forest age and forest connectivity on litter chemical attributes and fungal composition along the chronosequence of forest restoration. Forest age and forest connectivity were used as predictors, while environmental variables were defined as response variables and tested individually, and plot was included as a random factor. Model assumptions, including residual normality, errors homoscedasticity, collinearity, and absence of influential points and outliers were checked using the *performance* package (Lüdecke et al. 2021). If assumptions were violated, data were square-root transformed to meet model assumptions. We also used Spearman correlation analysis with the *corrplot* package to assess relationships among fungal guilds, tree diversity, and litter chemistry (*p* < 0.01).

To test our third hypothesis, a structural equation model (SEM) was constructed to assess the potential pathways through which forest age and forest connectivity influence tree diversity, litter biomass, litter chemical composition, and the abundance of fungal guilds. The conceptual model was developed based on the hypothesis that increasing forest age and connectivity may promote higher tree diversity, which in turn, affects litter chemistry and fungal guild composition (Figure S2, Table S2). This framework allows for the evaluation of direct relationships among forest age and connectivity, tree diversity, litter attributes, and fungal abundance, with indirect relationships being mediated by other variables. Specifically, we tested whether forest connectivity could indirectly affect fungal guild abundance by affecting tree diversity or by altering litter biomass and chemical composition as a function of restoration time. For that, we included restoration age and surrounding forest cover as predictors for forest age and forest connectivity, respectively.

Pastures were considered as a restoration starting point (0 years) and Primary forests were considered the endpoint (100 years). The first PCA axis was used as a proxy of litter chemistry composition, while the first NMDS axis was used as a proxy of litter fungal composition. Fungal guilds were included based on their relative abundance among classified fungal taxa.

We expected that older and more connected forests would correlate with higher tree diversity and greater leaf litter biomass. Increased tree diversity was expected to lead to more chemically diverse litter, influencing fungal guild composition by providing more heterogeneous substrates. Furthermore, forest age and connectivity were expected to affect fungal guild abundance through both direct and indirect pathways. Given that fungal guilds are influenced by vegetation, climate, and resource availability, we assumed autocorrelations between litter biomass and litter chemistry, and between the abundance of saprophytic and endophytic fungi. Therefore, these variables were thus included as correlated error in the SEM.

The SEM was tested using the *piecewiseSEM* package (Lefcheck 2016), which enables the evaluation of individual paths analysis. Each path was modeled using linear mixed-effects models, with plot included as a random effect, and the model assumptions were checked again with the *performance* package. The global model fit was assessed using the Fischer’s C statistics (*p* > 0.05), Akaike Information Criterion (AIC), and the goodness-of-fit index (GFI). For all models, standardized path coefficients (based on variable standard deviations) were calculated to allow comparison of the relative strength of each path within the SEM.

## 3. Results

### 3.1. Litter chemistry and fungal composition are affected by land use

We analyzed litter chemical attributes across different land use types by PCA and showed that the first two PCA axes accounted for 85.4% of the observed variation in litter characteristics (Figure 1A). The first axis separated primary and secondary forest litter from pasture litter. This separation was related to litter total N, S, and P content, which were higher in primary and secondary forests than in pasture. Pasture litter was also characterized by the lowest nutrient contents and highest δ^13^C values. The second axis separated the primary forest litter from that of the secondary forest and pasture. The primary forest had the highest δ^15^N values, and the secondary forest litter had the highest content of K and Zn (Figure 1A and Figure S3).

**Figure 1.**
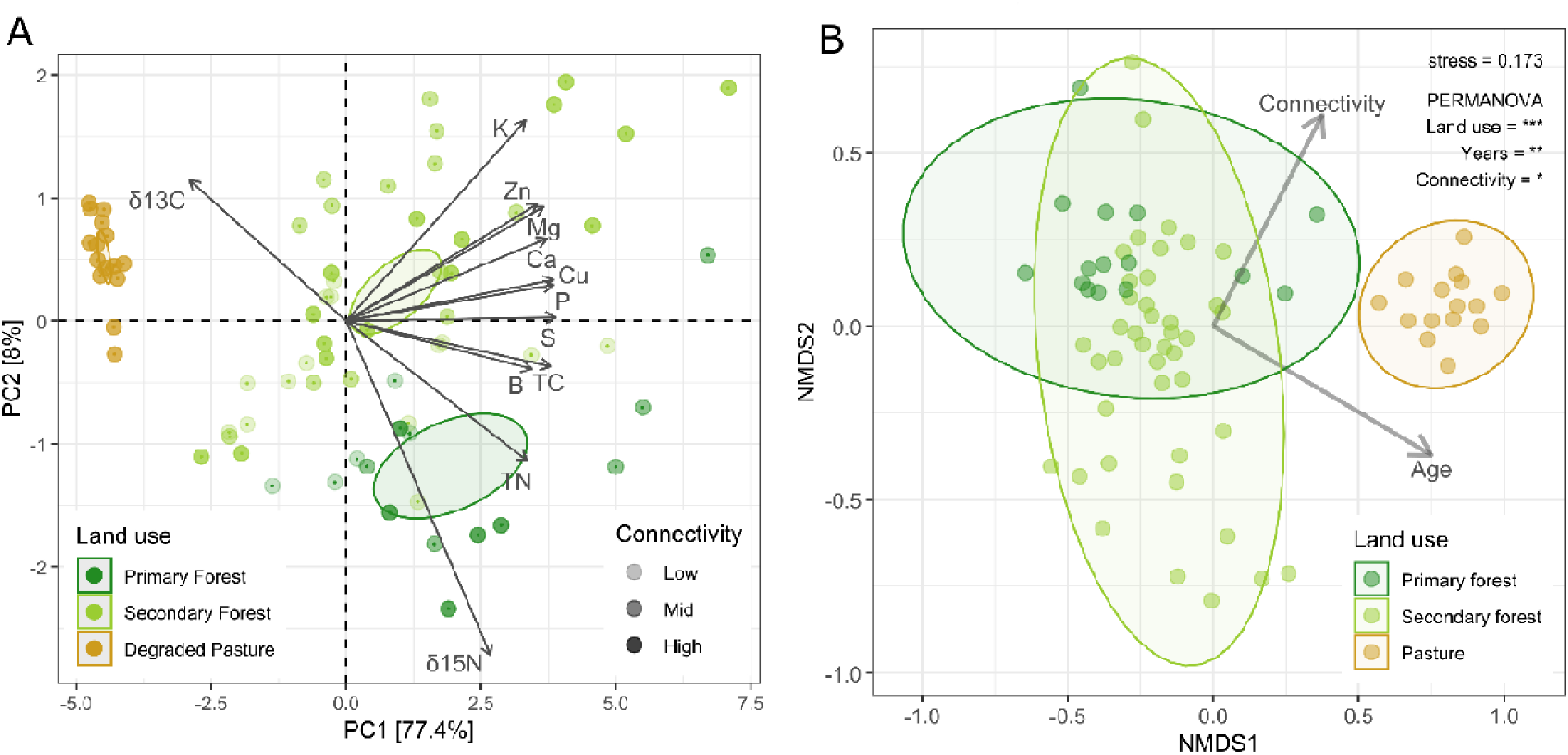
(A) The chemical composition of the litter layer under different land use types represented by a Principal Component Analysis (PCA). Ellipses show the confidence interval (*p* < 0.05). Litter fungal composition under different land-use types represented by a nonmetric multidimensional scaling (NMDS). Arrows indicate the effects of connectivity and age on fungal beta diversity. The ellipses represent the confidence intervals for land use types (p < 0.05).

We observed significant differences in fungal community composition across land uses (Figure 1B). Fungal composition of litter from both primary and secondary forests differed significantly from pastures (p < 0.001), whereas differences between the primary and secondary forest were smaller compared to differences from pasture litter. Forest age had a greater influence on fungal composition of litter than forest connectivity. Land use type (pasture vs. primary forest vs. secondary forest) explained 15.4% of the variation in fungal composition, while forest age and connectivity explained 4.4% and 3.2%, respectively. The fungal composition in secondary forests were highly heterogeneous, with little overlap in species abundance across samples. In contrast, fungal communities in pasture litter showed greater similarity, meaning they shared a more similar abundance of species across samples.

### 3.2. Fungal biomarkers and fungal guilds across pastures and forests

We identified a total of 2693 fungal ASVs, with an average of 87342 reads per sample. Two phyla dominated fungal community in the litter layer: Ascomycota, which was significantly more abundant in pasture litter compared to primary and secondary forests (Figure S3), and Basidiomycota, which was more abundant in primary forest litter, followed by secondary forest and pasture litter. Together, these two phyla represented over 90% of the total fungal community.

Using random forest models, we identified distinct fungal biomarkers associated with each litter type (Figure S4). In primary forest litter, the class Sordariomycetes (Ascomycota) was the most important and abundant biomarker. In contrast, Leotiomycetes (Ascomycota) and Rhizophlyctidomycetes (Chytridiomycota) were the most important biomarkers for secondary forest litter, despite their relatively low abundance. In pasture litter, the most important biomarkers was Ustilaginomycetes (Basidiomycota), while Dothideomycetes (Ascomycota) was the most abundant.

We also characterized the functional composition of fungal guilds across litter types (Figure S5). The most dominant guilds were, respectively, plant pathogens, wood saprotrophs, endophytes, and fungal parasites, which together comprised over 80% of the classified fungal guilds. Plant pathogens were more prevalent in primary forest litter, followed by secondary forest and pasture litter. Wood saprotrophs and endophytes were also more abundant in primary and secondary forests litter, than in pasture litter. In contrast, pasture litter presented higher abundance of soil saprotrophs, plant saprotrophs, and litter saprotrophs, compared to primary and secondary forests litter.

### 3.3. Both forest age and connectivity drives litter attributes and fungal guilds

Forest age and forest connectivity significantly explained the variation in litter chemistry, litter biomass, and fungal community composition (Figure 2; Table S4). Both forest age and connectivity were positively associated with the first principal component (PC1), which represented a gradient in litter chemistry. Litter chemistry was more strongly predicted by forest connectivity (β = 0.090, p = 0.010) than forest age (β = 0.040, p = 0.018). Conversely, litter biomass was significantly predicted by forest age (β = 0.650, p = 0.011), while the effect of forest connectivity was marginal (β = 1.060, p = 0.054). Fungal community composition, represented by the first axis of the NMDS, was significantly predicted by both forest age and connectivity, showing negative relationships (β = -0.005, p = 0.043; and β = -0.010, p = 0.042, respectively).

**Figure 2.**
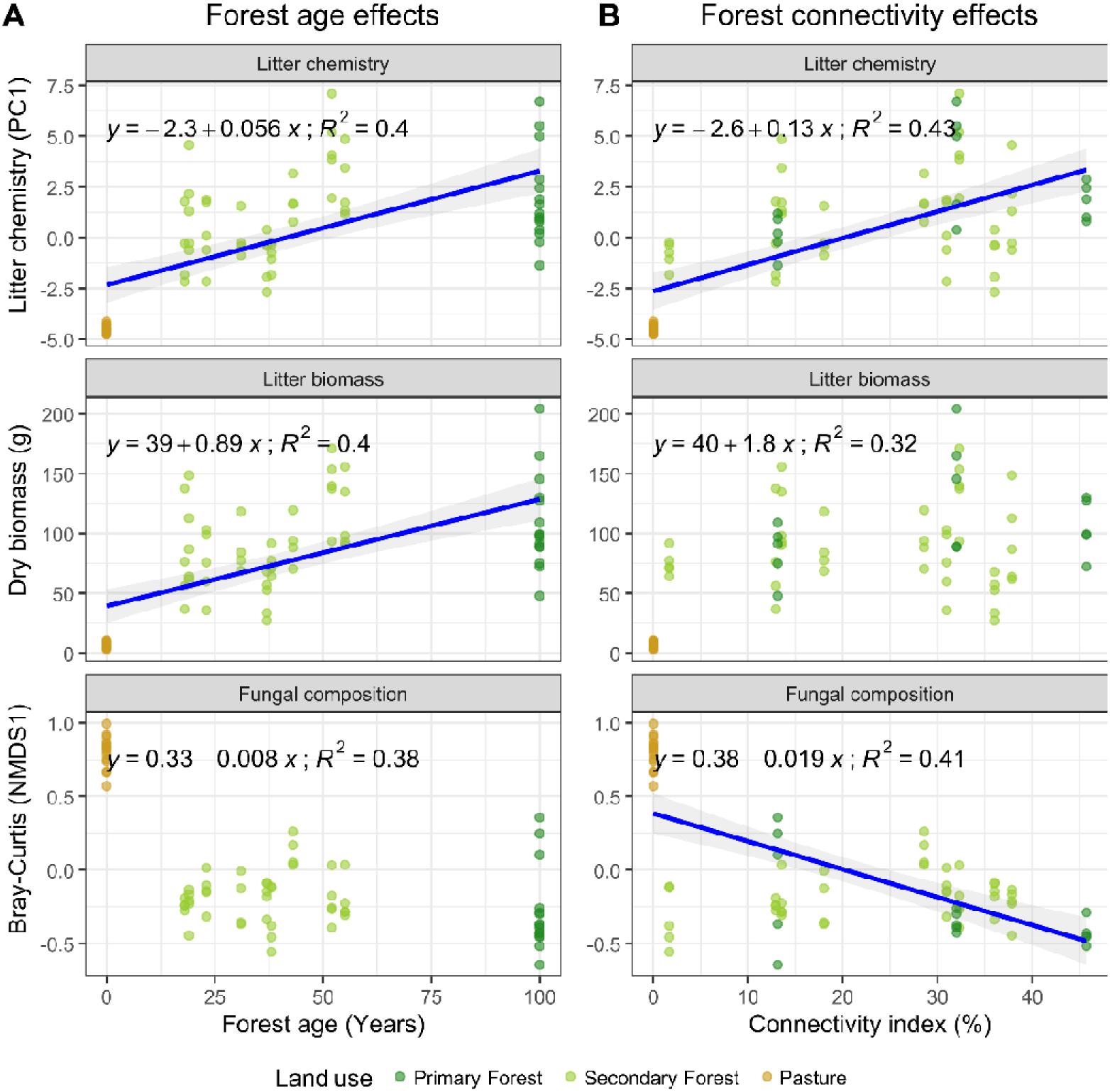
The effects of forest age (A) and forest connectivity (B) on litter chemistry, litter biomass and litter fungal composition. Litter chemistry is represented by the first axis of the PCA. Litter fungal composition is represented by the first axis of the NMDS based on the Bray-Curtis dissimilarity. Lines represents significant relationships between variables. R^2^ values represent the coefficient of correlation between variables.

Forest age also significantly influenced the abundance of specific fungal guilds in the litter layer (Figure 3; Table S4). Specifically, it was positively associated with endophytes (β = 0.030, p = 0.049) and had a marginal negative association with plant parasites (β = 0.0034, p = 0.062). Forest connectivity further shaped fungal functional composition (Figure 4; Table S4), showing significant positive effects on wood saprotrophs (β = 0.130, p = 0.002) and marginal effects on endophytes (β = 0.060, p = 0.075). In contrast, soil saprotrophs were negatively associated with forest connectivity (β = -0.0084, p = 0.002).

**Figure 3.**
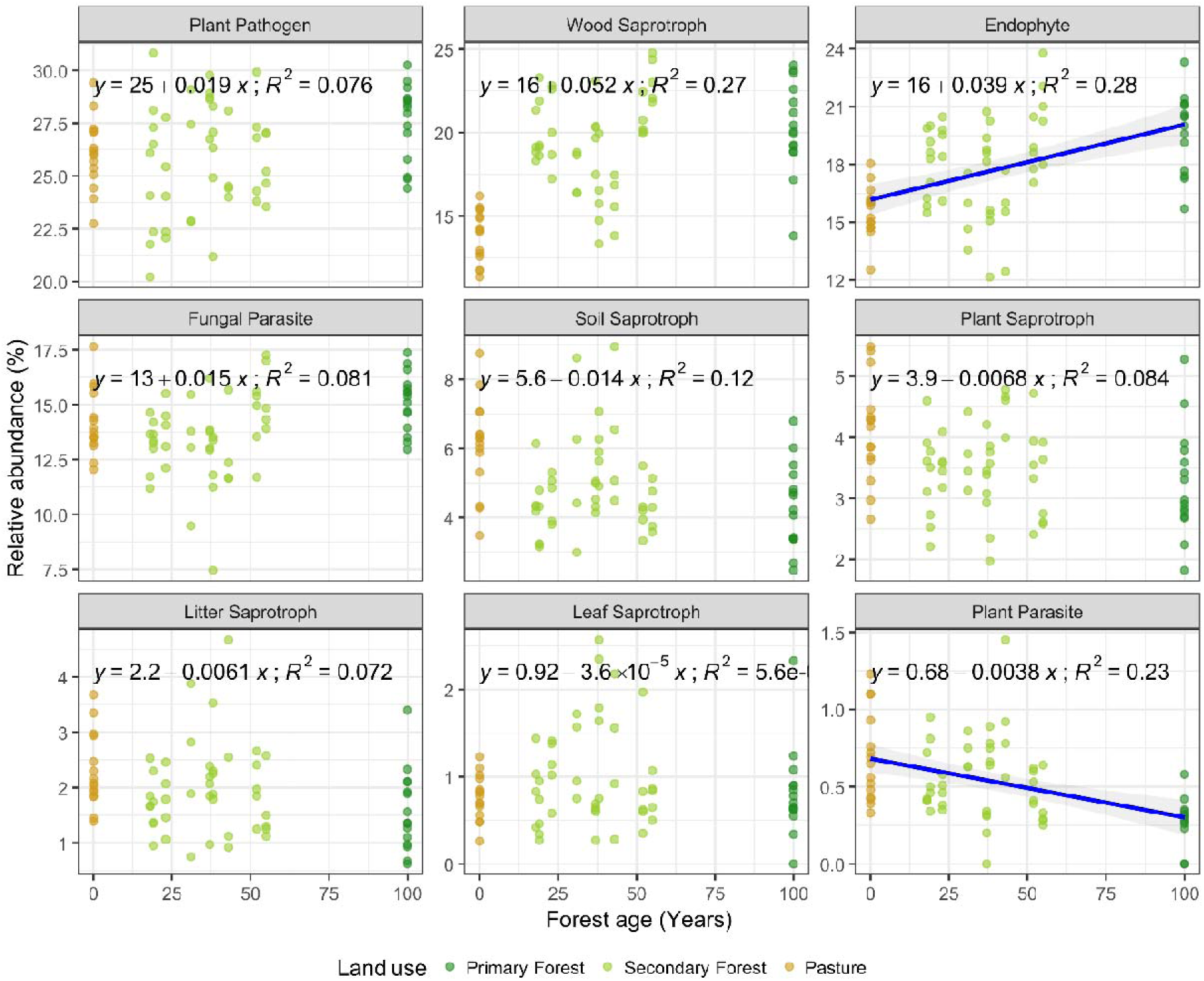
The effect of forest age on the relative abundance of fungal guilds during forest restoration. Lines represents significant relationships between fungal guilds and forest age and forest connectivity. R^2^ values represent the coefficient of correlation between variables.

**Figure 4.**
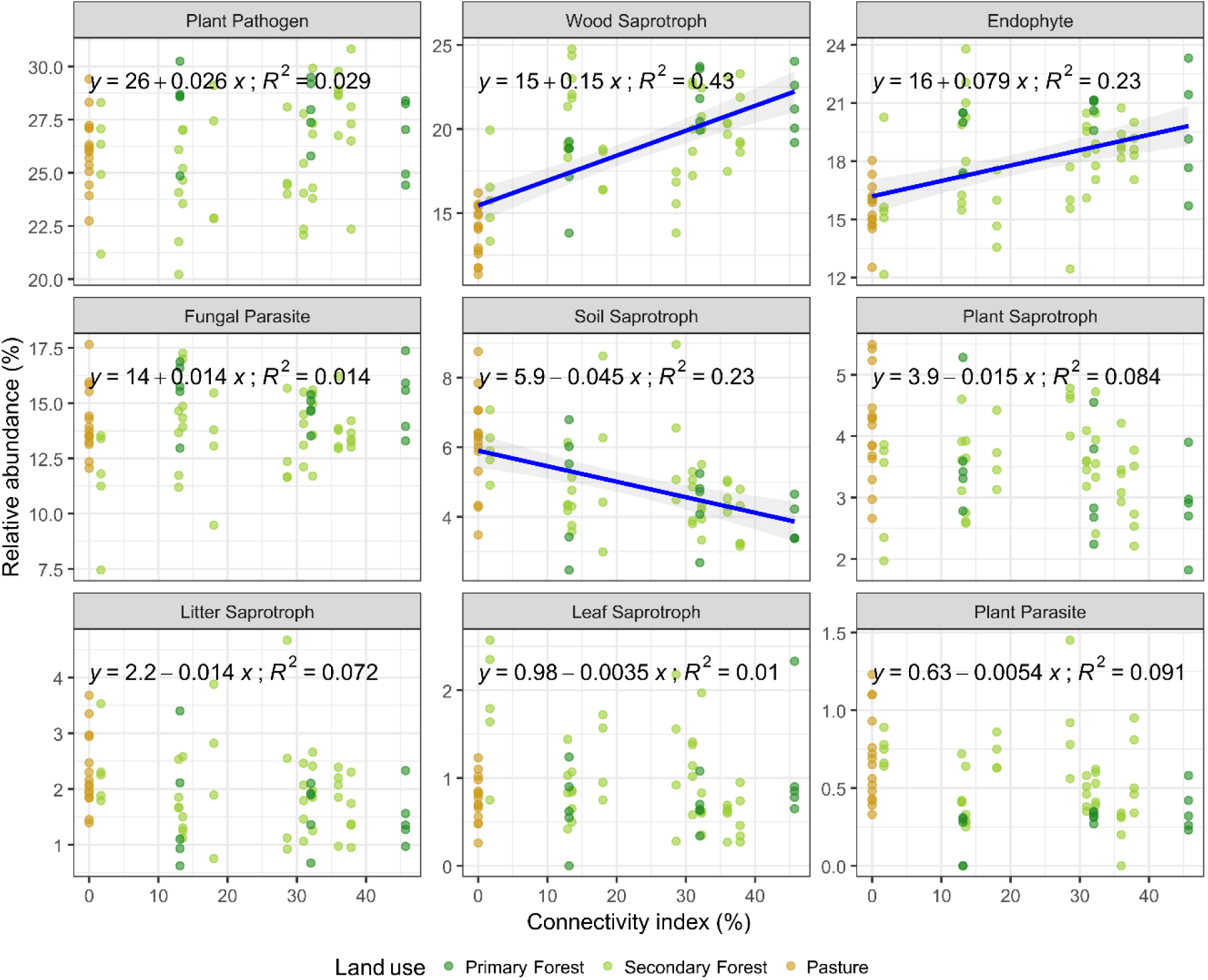
The effect of forest connectivity on the relative abundance of fungal guilds during forest restoration. Lines represents the relationships between fungal guilds and forest age and forest connectivity. R^2^ values represent the coefficient of correlation between variables.

### 3.4. Environmental drivers of fungal guilds

Fungal guilds in leaf litter were significantly correlated with tree diversity, litter biomass, and nutrient content (Figure S6). Wood saprotrophs and endophytes showed positive correlations with most environmental variables, except with δ^13^C. In contrast, plant saprotrophs, soil saprotrophs, litter saprotrophs, and plant parasites were negatively correlated with these environmental variables. The strongest positive correlations were observed between wood saprotrophs and litter δ^15^N (ρ = 0.74), and between endophytes and litter δ^15^N (ρ = 0.72). Additionally, litter P and tree diversity were strongly correlated with wood saprotrophs abundance (both ρ = 0.65), while litter S and litter biomass were strongly associated with endophyte abundance (ρ = 0.53 and 0.50, respectively). Further, the strongest negative correlations included plant parasites and δ^15^N (ρ = -0.68), as well as soil saprotrophs and tree diversity (ρ = -0.44).

Structural equation modeling (SEM) showed that tree diversity was more strongly affected by forest connectivity (standardized path coefficient = 0.49) than by forest age (0.35) (Figure 5; Table S5). Tree diversity, in turn, had significant positive effects on both litter biomass (0.64) and litter chemistry (0.56), whereas forest age did not show a significant direct effect on either variable. Regarding fungal guilds, wood saprotrophs were positively affected by both forest connectivity (0.50) and litter chemistry (0.27). Endophyte abundance was marginally affected by forest connectivity (0.37) but not significantly influenced by forest age (0.31).

**Figure 5.**
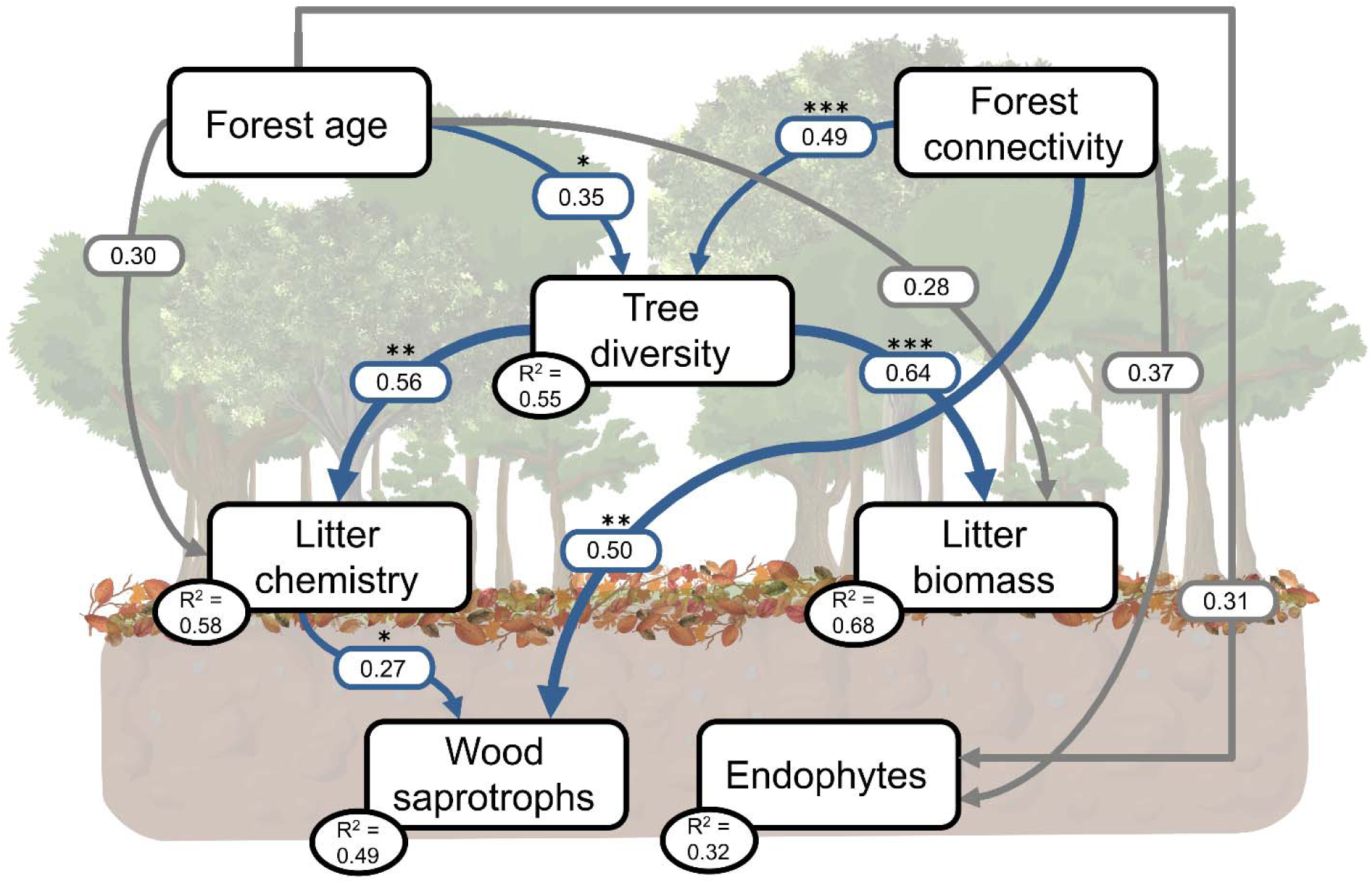
Structural Equation Model (SEM) shows the effects of forest age and connectivity during restoration on litter biomass, litter chemistry and litter fungal guilds. Arrows show the standardized path coefficient of linear regressions presented in Table S2. Blue arrow represent significant and positive relationships. Grey arrow represent non-significant relationships. Asterisks indicate the significance level of each path (* = *p* < 0.05; ** = *p* < 0.01; and *** = *p* < 0.001). The SEM fit the data well, represented by the global goodness-of-fit with Fischer’s C = 12.803; *p* = 0.687; *df* = 16; *n* = 75. The variation explained in each variable (R^2^) represents the marginal response and is shown inside each box.

## 4. Discussion

In this study, we investigated the influence of forest age and connectivity on the fungal composition and functioning in forest litter during the restoration of Atlantic Forest fragments in Brazil. Our findings revealed that forest connectivity has moderate yet significant effects on fungal composition and litter chemistry. Specifically, forest connectivity accelerates the functional recovery of microbial decomposers by enhancing tree diversity and enriching the litter layer, strengthening nutrient cycling and supporting forest restoration. These connections provide valuable insights into how litter microbes help restore ecosystem functions in lands previously degraded by grazing, from compacted, nutrient-poor, and nearly bare soils to healthy forested ecosystems.

### 4.1. How does forest age and connectivity affect litter attributes and fungal composition?

In our study, both forest connectivity and forest age correlated well with litter chemistry. However, only forest age had a significant effect on litter biomass. This suggests that restored tropical forests progressively enhance tree leaf biomass over time, resulting in more litterfall and an accumulation of litter biomass (Souza et al. 2019). In contrast, older and connected secondary forests typically have higher tree diversity compared to older and isolated secondary forests (Rother et al., 2023; Zucchi et al., 2018), which affects the overall composition of litter chemistry and accumulation of litter biomass. This pattern likely occurs because forests with high tree diversity produce litter with wider variety of complex carbon compounds, which in turn, shapes the fungal community by promoting mutualistic interactions that enhance decomposition efficiency (García-Palacios et al. 2013; Pei et al. 2017). In the context of forest restoration, a higher percentage of surrounding forest cover promotes species conservation and increases the quality of natural regeneration (Chazdon et al. 2025). Furthermore, greater connectivity may accelerate litter decomposition and increase nutrient release into the soil because diverse microbial communities are more efficient at breaking down organic matter than simplified ones (Buresova et al. 2019). Although we did not use litter bags to measure the decomposition process over time, we observed an increase in many fungal taxa related to litter degradation.

Additionally, our results showed that the Fabaceae family was the most abundant and diverse in secondary forests (Table S5), suggesting that a significant portion of litter N and P may originate from these species. For example, Fabaceae trees are fast-growing and capable of biological N fixation, producing high-protein litter with elevated N and P concentrations (Neves et al. 2022; Wang et al. 2022). We also found that litter N and P content were positively correlated with fungal guilds such as endophytes and wood saprotrophs. This indicates that litter nutrient availability is a key driver of fungal community shifts, altering the predominance of oligotrophic Ascomycota to copiotrophic Basidiomycota (Guo et al. 2024). This microbial transition is largely induced by changes in tree species composition (Urbanová et al. 2015) that further affect environmental factors such as soil moisture, pH, and nutrient availability (Huang et al. 2023).

We also observed a shift from oligotrophic to copiotrophic microorganisms in our study. Pasture litter was dominated by Ascomycota and had a low abundance of Basidiomycota, whereas primary forest litter showed the opposite trend, with secondary forests showing intermediate values (Figure S3). These transitions likely reflect changes in litter quality and decomposition stages during forest succession. In the early successional stages, litter is typically rich in cellulose but poor in nutrients, favoring Ascomycota. In contrast, more mature forests produce litter richer in lignin and nutrients, which supports Basidiomycota (Baldrian 2017; Hannula et al. 2017). This Ascomycota-to-Basidiomycota ratio is a strong indicator of forest successional stage (Gourmelon et al. 2016). In our case, forest connectivity was more strongly associated with litter chemistry and fungal composition than with forest age, suggesting that higher percentages of forest cover enhance nutrient concentration in litter, further shaping the fungal composition. Together, these effects alters the fungal community from oligotrophic to copiotrophic taxa, which potentially may lead to faster decomposition and nutrient cycling. We conclude that forest connectivity enhanced the shift from oligotrophic to copiotrophic fungi, likely reflecting a better resto of litter decomposition processes.

### 4.2. Forest connectivity and its relationship with saprophytic and endophytic communities

It is well established that increasing connectivity between forest fragments enhances the diversity and functionality of plants and macrofauna during forest restoration (Rother et al. 2018; Matos et al. 2020). In this study, we demonstrate that the effects of forest age and connectivity also occurs at smaller scales, particularly in litter chemistry and fungal communities. These effects become more pronounced at intermediate and late stages of restoration compared to the earliest stages. While fungal composition is directly linked to litter chemistry and host specificity (Zhang et al. 2020), tree composition is more strongly associated with broader environmental variables such as forest connectivity and regional edaphic conditions (Zucchi et al. 2018). This creates an indirect pathway of forest connectivity influencing fungal composition in the litter layer.

Endophytic and pathogenic fungi in fresh leaves depend on carbon supplied by the host plant (Vincent et al. 2016; Zhang et al. 2020), but is suggested that many groups continue their life cycle in the leaf litter after senescence (Otsing et al. 2018). Some studies further suggests that endophytes remain an active and substantial part of the litter fungal community even one year after litterfall (Guerreiro et al. 2018). At the same time, many endophytic fungi shift to a saprophytic lifestyle, becoming part of the initial decomposer community and influencing the succession of later fungal groups adapted to degrading more recalcitrant compounds, such as lignin (Kirker et al. 2020). Here, endophytic fungi and litter biomass were mainly associated with forest age, suggesting that litter accumulation in older forests is more related to fresh litterfall. In contrast, saprotrophs and litter chemistry were more strongly associated with forest connectivity, implying an association with older, more decomposed litter characterized by higher C/N ratios and greater lignin content (Leifheit et al. 2024). This interpretation is further supported by the observed shift in the Ascomycota-to-Basidiomycota ratio in our litter samples, with Basidiomycota becoming more dominant in later stages of litter decomposition.

Specifically, Sordariomycetes (Ascomycota) and Agaricomycetes (Basidiomycota) are important groups of decomposers of complex substrates such as lignin and tannin (Veen et al. 2021), and were more abundant in our old-growth forests. In contrast, Dothideomycetes (Ascomycota) was a biomarker for pasture litter, previously shown to be strongly associated with higher cellulose content in litter (Veen et al. 2021). Interestingly, both Sordariomycetes and Dothideomycetes are cosmopolitan and highly versatile, capable of functioning as either saprophytes or endophytes, depending on resource availability (Freitas et al. 2021). These functional adaptations contribute to fungal multifunctionality and niche specialization in the litter layer (Bastida et al. 2016; Bhatnagar et al. 2018).

Furthermore, litter isotope signature (δ^13^C and δ^15^N) was significantly correlated with both forest age and forest connectivity, as well as with most fungal guilds with in the litter. On one hand, saprophytes and endophytes were associated with higher δ^15^N values, while plant parasites were associated with lower δ^13^C values. This suggests that microbes preferentially consume lighter isotopes (^12^C and ^14^N) during decomposition, resulting in an enrichment of heavier isotopes in the remaining litter (Bieluczyk, et al., 2023a). δ^15^N enrichment is often attributed to both microbial biomass incorporation into decomposing litter (Bieluczyk, et al., 2023b), and increased biological nitrogen fixation (Zanini et al. 2021). On the other hand, the relationships between litter isotopic signature and forest age connectivity are likely influenced by the origin and decomposition stage of organic matter. For instance, higher δ^13^C values are often derived from C_4_ plants and are indicative of early-stages restoration (Zanini et al. 2021), but they can also reflect older, more recalcitrant litter, as heavier isotopes form stronger chemical bonds that are more resistant to microbial breakdown, resulting in higher δ^13^C (Xiong et al. 2020). Although no strong evidence directly links forest connectivity to litter isotope signature, we speculate that the enrichment of δ^15^N in more connected forests may result from both the presence of nitrogen-fixing bacteria, which supply additional N to other decomposer microorganisms, and by the accumulation of δ^15^N in recalcitrant organic matter.

## 5. Conclusions

Increased forest connectivity led to higher proportions of wood saprotrophs and endophytes in the litter layer, though this connectivity effect was more evident in the intermediate and late stages of forest succession. Secondary forests with at least 30% connectivity were high in wood saprotrophs and endophytes in the litter layer, and these proportions increased along the chronosequence of forest restoration as connectivity increased. Our findings suggest that enhancing forest connectivity to 30% is a promising measure for restoring the fungal litter community, especially in intermediate and late stages of restoration. Although intermediate and late restoration stages had higher abundance of these fungal guilds compared to early secondary forests, enhancing connectivity can benefit all stages of restoration.

The litter layer responds well to both forest age and connectivity, making it a useful metric for assessing ecosystem services in secondary forests, such as promoting fungal diversity and supporting carbon sequestration and storage through microbial activity. Whether a loss of forest connectivity leads to declines in fungal communities and reduced nutrient cycling, potentially contributing to forest degradation through proliferation of more plant parasites and pathogens, requires further studies. Investigating these effects is relevant, as it may explain how climate change effects may exacerbate future declines in plant diversity and alter community composition. Our study shows that these processes may, at least in part, be reversed by restoring forests while considering spatial connectivity of the new and existing forests areas. While the practical implications of increasing connectivity on microbial attributes may seem minor, these changes can increase the resilience of newly restored forests in the face of future climate change scenarios when considered at a broader scale and over the long term. The One Health concept recognizes that human, animal, and plant health are closely linked to environmental and ecosystem stability.

## Acknowledgments

The authors would like to thank the collaboration of the São Paulo Research Foundation (FAPESP) and the Dutch Research Council (NWO) together for financial support in the project FAPESP/NWO – BioFor (2018/19000-4) and the project FAPESP/NWO – ReSeeD (2018/19000-6). GLM thanks FAPESP for the PhD scholarship (Process 2022/05561-0) and the Coordenação de Aperfeiçoamento de Pessoal de Nível Superior - Brasil (CAPES) for granting the international scholarship CAPES-PrInt (Process 88887.831610/2023-00). DZ was supported by a Horizon Europe Marie Skłodowska-Curie Postdoctoral Fellowship (grant agreement number 101110604 – BelowGround). WB thanks FAPESP for funding (Process 2021/00976-4) and the post-doctoral scholarship (Process #2023/18333-8). All authors thank Prof. Pedro Brancalion and the FAPESP-NWO NewFor project (2018/18416-2) for providing the database of the plant composition of the restoration areas.

## Author Contributions

**Guilherme Lucio Martins**: conceptualization, formal analysis, investigation, methodology, visualization, writing - original draft; **Dina in ’t Zandt**: conceptualization, formal analysis, supervision, writing - review & editing; **Gabriel Silvestre Rocha**: conceptualization, investigation, validation, writing - review & editing; **Luis Fernando Merloti**: conceptualization, investigation, data curation, writing - review & editing; **Wanderlei Bieluczyk**: conceptualization; investigation, validation, writing - review & editing; **Robert Timmers**: conceptualization, data curation, writing - review & editing; **Ricardo Ribeiro Rodrigues**: data curation, writing - review & editing; **Siu Mui Tsai**: conceptualization, funding acquisition, project administration, resources, supervision, writing - review & editing; **Wim H. van der Putten**: conceptualization, funding acquisition, resources, supervision, writing - review & editing.

## References

Alvares CA, Stape JL, Sentelhas PC, et al (2013) Köppen’s climate classification map for Brazil. Meteorologische Zeitschrift 22:711–728. 10.1127/0941-2948/2013/0507

Baldrian P (2017) Forest microbiome: Diversity, complexity and dynamics. FEMS Microbiol Rev 41:109–130. 10.1093/femsre/fuw040

Bardgett RD, Van Der Putten WH (2014) Belowground biodiversity and ecosystem functioning. Nature 515:505–511. 10.1038/nature13855

Bastida F, Torres IF, Moreno JL, et al (2016) The active microbial diversity drives ecosystem multifunctionality and is physiologically related to carbon availability in Mediterranean semi-arid soils. Mol Ecol 25:4660–4673. 10.1111/mec.13783

Bates D, Mächler M, Bolker BM, Walker SC (2015) Fitting linear mixed-effects models using lme4. J Stat Softw 67:. 10.18637/jss.v067.i01

Bhatnagar JM, Peay KG, Treseder KK (2018) Litter chemistry influences decomposition through activity of specific microbial functional guilds. Ecol Monogr 88:429–444. 10.1002/ecm.1303

Bieluczyk W, Asselta FO, Navroski D, et al (2023a) Linking above and belowground carbon sequestration, soil organic matter properties, and soil health in Brazilian Atlantic Forest restoration. J Environ Manage 344:. 10.1016/j.jenvman.2023.118573

Bieluczyk W, Merloti LF, Cherubin MR, et al (2023b) Forest restoration rehabilitates soil multifunctionality in riparian zones of sugarcane production landscapes. Science of the Total Environment 888:. 10.1016/j.scitotenv.2023.164175

Brancalion PHS, Schweizer D, Gaudare U, et al (2016) Balancing economic costs and ecological outcomes of passive and active restoration in agricultural landscapes: the case of Brazil. Biotropica 48:856–867. 10.1111/btp.12383

Buresova A, Kopecky J, Hrdinkova V, et al (2019) Succession of Microbial Decomposers Is Determined by Litter Type, but Site Conditions Drive Decomposition Rates

Callahan BJ, McMurdie PJ, Rosen MJ, et al (2016) DADA2: High-resolution sample inference from Illumina amplicon data. Nat Methods 13:581–583. 10.1038/nmeth.3869

Carlucci MB, Brancalion PHS, Rodrigues RR, et al (2020) Functional traits and ecosystem services in ecological restoration. Restor Ecol 28:1372–1383

César RG, Moreno V de S, Coletta GD, et al (2021) It is not just about time: Agricultural practices and surrounding forest cover affect secondary forest recovery in agricultural landscapes. Biotropica 53:496–508. 10.1111/btp.12893

Chai Y, Cao Y, Yue M, et al (2019) Soil abiotic properties and plant functional traits mediate associations between soil microbial and plant communities during a secondary forest succession on the Loess Plateau. Front Microbiol 10:. 10.3389/fmicb.2019.00895

Chazdon RL, Blüthgen N, Brancalion PHS, et al (2025) Drivers and benefits of natural regeneration in tropical forests. Nature Reviews Biodiversity 1:298–314. 10.1038/s44358-025-00043-y

Chen C, Bongers FJ, Schmid B, et al (2025) Ecosystem consequences of functional diversity in forests and implications for restoration. New Phytologist

de Lima RAF, Oliveira AA, Pitta GR, et al (2020) The erosion of biodiversity and biomass in the Atlantic Forest biodiversity hotspot. Nat Commun 11:. 10.1038/s41467-020-20217-w

England LS, Vincent ML, Trevors JT, Holmes SB (2004) Extraction, detection and persistence of extracellular DNA in forest litter microcosms. Mol Cell Probes 18:313–319. 10.1016/j.mcp.2004.05.001

Fahrig L (2013) Rethinking patch size and isolation effects: The habitat amount hypothesis. J Biogeogr 40:1649–1663. 10.1111/jbi.12130

Farquhar GD, O’Leary MH, Berry JA (1982) On the relationship between carbon isotope discrimination and the intercellular carbon dioxide concentration in leaves. Aust J Plant Physiol 9:121–137. 10.1071/PP9820121

Freitas MLR, Gomes AAM, Rosado AWC, Pereira OL (2021) Cladosporium species from submerged decayed leaves in Brazil, including a new species and new records. Phytotaxa 482:223–239

García-Palacios P, Maestre FT, Kattge J, Wall DH (2013) Climate and litter quality differently modulate the effects of soil fauna on litter decomposition across biomes. Ecol Lett 16:1045–1053. 10.1111/ele.12137

Gourmelon V, Maggia L, Powell JR, et al (2016) Environmental and geographical factors structure soil microbial diversity in new caledonian ultramafic substrates: A metagenomic approach. PLoS One 11:. 10.1371/journal.pone.0167405

Guerreiro MA, Brachmann A, Begerow D, Peršoh D (2018) Transient leaf endophytes are the most active fungi in 1-year-old beech leaf litter. Fungal Divers 89:237–251. 10.1007/s13225-017-0390-4

Guo X, Wang S, Wang C, et al (2024) The Changes, Aggregation Processes, and Driving Factors for Soil Fungal Communities during Tropical Forest Restoration. Journal of Fungi 10:. 10.3390/jof10010027

Hannula SE, Morriën E, De Hollander M, et al (2017) Shifts in rhizosphere fungal community during secondary succession following abandonment from agriculture. ISME Journal 11:2294–2304. 10.1038/ismej.2017.90

Hatfield JH, Banks-Leite C, Barlow J, et al (2024) Constraints on avian seed dispersal reduce potential for resilience in degraded tropical forests. Funct Ecol 38:315–326. 10.1111/1365-2435.14471

Huang K, Guo Z, Zhao W, et al (2023) Response of fungal communities to afforestation and its indication for forest restoration. For Ecosyst 10:. 10.1016/j.fecs.2023.100125

Kirker GT, Bishell A, Cappellazzi J, et al (2020) Role of leaf litter in above-ground wood decay. Microorganisms 8:. 10.3390/microorganisms8050696

Lefcheck JS (2016) piecewiseSEM: Piecewise structural equation modelling in r for ecology, evolution, and systematics. Methods Ecol Evol 7:573–579. 10.1111/2041-210X.12512

Leifheit EF, Camenzind T, Lehmann A, et al (2024) Fungal traits help to understand the decomposition of simple and complex plant litter. FEMS Microbiol Ecol 100:. 10.1093/femsec/fiae033

Liu C, Cui Y, Li X, Yao M (2021) Microeco: An R package for data mining in microbial community ecology. FEMS Microbiol Ecol 97:. 10.1093/femsec/fiaa255

Liu S, Plaza C, Ochoa-Hueso R, et al (2023) Litter and soil biodiversity jointly drive ecosystem functions. Glob Chang Biol 29:6276–6285. 10.1111/gcb.16913

Lüdecke D, Ben-Shachar M, Patil I, et al (2021) performance: An R Package for Assessment, Comparison and Testing of Statistical Models. J Open Source Softw 6:3139. 10.21105/joss.03139

Malavolta E, Vitti GC, de Oliveira SA (1989) Evaluation of the nutritional state of plants: principles and applications. Associação Brasileira para Pesquisa da Potasse e do Fosfato

Matos FAR, Magnago LFS, Aquila Chan Miranda C, et al (2020) Secondary forest fragments offer important carbon and biodiversity cobenefits. Glob Chang Biol 26:509–522. 10.1111/gcb.14824

Neves NM, Paula RR, Araujo EA, et al (2022) Contribution of legume and non-legume trees to litter dynamics and C-N-P inputs in a secondary seasonally dry tropical forest. IForest 15:8–15. 10.3832/ifor3442-014

Nguyen NH, Song Z, Bates ST, et al (2016) FUNGuild: An open annotation tool for parsing fungal community datasets by ecological guild. Fungal Ecol 20:241–248. 10.1016/j.funeco.2015.06.006

Nilsson RH, Larsson KH, Taylor AFS, et al (2019) The UNITE database for molecular identification of fungi: Handling dark taxa and parallel taxonomic classifications. Nucleic Acids Res 47:D259–D264. 10.1093/nar/gky1022

Otsing E, Barantal S, Anslan S, et al (2018) Litter species richness and composition effects on fungal richness and community structure in decomposing foliar and root litter. Soil Biol Biochem 125:328–339. 10.1016/j.soilbio.2018.08.006

Oksanen, J., Blanchet, F.G., Friendly, M., Kindt, R., Legendre, P., Mcglinn, D., Minchin, P.R., O’hara, R.B., Simpson, G.L., Solymos, P., et al. (2019). Package “vegan” Community Ecology Package.

Pei Z, Leppert KN, Eichenberg D, et al (2017) Leaf litter diversity alters microbial activity, microbial abundances, and nutrient cycling in a subtropical forest ecosystem. Biogeochemistry 134:163–181. 10.1007/s10533-017-0353-6

Poorter L, Craven D, Jakovac CC, et al (2021) Multidimensional tropical forest recovery. Science (1979) 374:1370–1376. 10.1126/science.abh362

Rezende CL, Scarano FR, Assad ED, et al (2018) From hotspot to hopespot: An opportunity for the Brazilian Atlantic Forest. Perspect Ecol Conserv 16:208–214. 10.1016/j.pecon.2018.10.002

Rodrigues RR, Gandolfi S, Nave AG, et al (2011) Large-scale ecological restoration of high-diversity tropical forests in SE Brazil. For Ecol Manage 261:1605–1613. 10.1016/j.foreco.2010.07.005

Rodrigues RR, Lima RAF, Gandolfi S, Nave AG (2009) On the restoration of high diversity forests: 30 years of experience in the Brazilian Atlantic Forest. Biol Conserv 142:1242– 1251. 10.1016/j.biocon.2008.12.008

Rother DC, Romanelli JP, Rodrigues RR (2023) Historical trajectory of restoration practice and science across the Brazilian Atlantic Forest. Restor Ecol 31

Rother DC, Vidal CY, Fagundes IC, et al (2018) How Legal-Oriented Restoration Programs Enhance Landscape Connectivity? Insights From the Brazilian Atlantic Forest. Trop Conserv Sci 11:. 10.1177/1940082918785076

Salviano IR, Gardon FR, dos Santos RF (2021) Ecological corridors and landscape planning: a model to select priority areas for connectivity maintenance. Landsc Ecol 36:3311–3328. 10.1007/s10980-021-01305-8

Saura S, Bastin L, Battistella L, et al (2017) Protected areas in the world’s ecoregions: How well connected are they? Ecol Indic 76:144–158. 10.1016/j.ecolind.2016.12.047

Souza SR, Veloso MDM, Espírito-Santo MM, et al (2019) Litterfall dynamics along a successional gradient in a Brazilian tropical dry forest. For Ecosyst 6:. 10.1186/s40663-019-0194-y

Timmers R, Côrtes MC, van Kuijk M, et al (2025) Landscape-scale forest cover shapes the complexity of seed-dispersal networks in regenerating forest fragments. Biol Conserv 309:. 10.1016/j.biocon.2025.111312

Toju H, Tanabe AS, Yamamoto S, Sato H (2012) High-coverage ITS primers for the DNA-based identification of ascomycetes and basidiomycetes in environmental samples. PLoS One 7:. 10.1371/journal.pone.0040863

Urbanová M, Šnajdr J, Baldrian P (2015) Composition of fungal and bacterial communities in forest litter and soil is largely determined by dominant trees. Soil Biol Biochem 84:53–64. 10.1016/j.soilbio.2015.02.011

Veen GF, ten Hooven FC, Weser C, Hannula SE (2021) Steering the soil microbiome by repeated litter addition. Journal of Ecology 109:2499–2513. 10.1111/1365-2745.13662

Vieira SA, Alves LF, Duarte-Neto PJ, et al (2011) Stocks of carbon and nitrogen and partitioning between above-and belowground pools in the Brazilian coastal Atlantic Forest elevation range. Ecol Evol 1:421–434. 10.1002/ece3.41

Vincent JB, Weiblen GD, May G (2016) Host associations and beta diversity of fungal endophyte communities in New Guinea rainforest trees. Mol Ecol 25:825–841. 10.1111/mec.13510

Wang S, Zhao S, Yang B, et al (2022) The carbon and nitrogen stoichiometry in litter-soil-microbe continuum rather than plant diversity primarily shapes the changes in bacterial communities along a tropical forest restoration chronosequence. Catena (Amst) 213:. 10.1016/j.catena.2022.106202

Wang, C., and Kuzyakov, Y. (2024). Soil organic matter priming: The pH effects. Global Change Biology 30, e17349. 10.1111/gcb.17349

Xiong X, Zhang H, Deng Q, et al (2020) Soil organic carbon turnover following forest restoration in south China: Evidence from stable carbon isotopes. For Ecol Manage 462:. 10.1016/j.foreco.2020.117988

Zanini AM, Mayrinck RC, Vieira SA, et al (2021) The effect of ecological restoration methods on carbon stocks in the Brazilian Atlantic Forest. For Ecol Manage 481:. 10.1016/j.foreco.2020.118734

Zhang N, Bruelheide H, Li Y, et al (2020) Community and neighbourhood tree species richness effects on fungal species in leaf litter. Fungal Ecol 47:. 10.1016/j.funeco.2020.100961

Zhang X, Wang L, Zhou W, et al (2022) Changes in litter traits induced by vegetation restoration accelerate litter decomposition in Robinia pseudoacacia plantations. Land Degrad Dev 33:179–192. 10.1002/ldr.4136

Zucchi MI, Sujii PS, Mori GM, et al (2018) Genetic diversity of reintroduced tree populations in restoration plantations of the Brazilian Atlantic Forest. Restor Ecol 26:694–701. 10.1111/rec.12620

